# *BioCell2XML*: A novel tool for converting cell lineage data from SIMI BioCell to MaMuT (Fiji)

**DOI:** 10.1101/554873

**Authors:** Markus Pennerstorfer, Günther Loose, Carsten Wolff

**Affiliations:** Humboldt-Universität zu Berlin, Institut für Biologie, Vergleichende Zoologie, Phillippstr. 13 Haus 2, 10115, Berlin

**Author notes:** Günther Loose.

## Abstract

Computer-assisted 4D manual cell tracking has been a valuable method for understanding spatial-temporal dynamics of embryogenesis (e.g., Stach & Anselmi, 2015; Vellutini et al., 2017; Wolff et al., 2018) since the method was introduced in the late 1990s. Since two decades *SIMI^®^ BioCell* (Schnabel et al., 1997), a software which initially was developed for analyzing data coming from the, at that time new technique of 4D microscopy, is in use. Many laboratories around the world use *SIMI BioCell* for the manual tracing of cells in embryonic development of various species to reconstruct cell genealogies with high precision. However, the software has several disadvantages: Limits in handling very large data sets, the virtually no maintenance over the last ten years (bound to older Windows versions), the difficulty to access the created cell lineage data for analyses outside *SIMI BioCell*, and the high cost of the program. Recently, bioinformatics, in close collaboration with biologists, developed new lineaging tools that are freely available through the open source image processing platform *Fiji*.

Here we introduce a software tool that allows conversion of *SIMI BioCell* lineage data to a format that is compatible with the *Fiji* plugin *MaMuT* (Wolff et al., 2018). Hereby we intend to maintain the usability of *SIMI BioCell* created cell lineage data for the future and, for investigators who wish to do so, facilitate the transition from this software to a more convenient program.

## Introduction

4D cell lineage reconstruction is performed on multifocal time-lapse videos of developing embryos or cultured cells. The image data is recorded with a specific microscopic setup (Schnabel et al., 1997), consisting of a microscope, camera(s), a motorized stage, and a controlling software. Today, all major manufacturers of research microscopes provide such equipment and it is used commonly in many developmental biology and cell biology laboratories around the world. The lineage data constitutes cell positions in time and space, which are saved as x/y/z/t coordinates and created by manually placing spots in the images at the respective locations (e.g. the nuclei) and time frames, and the genealogical relationships between these spots, which are saved in different ways depending on the program used.

*SIMI BioCell* is a software designed for manually tracking cells in developing embryos. It requires the use of a template, a prepared file with a predefined lineage structure. Cell names, numbers of divisions, or division times are defined in the template, but modified as the lineage data is entered. Users can choose between different templates, e.g. containing standard lineage trees of *C. elegans* at different temperatures. Additional data can be entered: Custom spot name or cell name, spot size, spot color, spot shape, cell fate, and comments. Cellular or acellular objects that are not part of the embryo’s cell set (e.g. polar bodies, nurse cell remnants, germ plasm, etc.) are difficult to include without “cheating” around the template’s structure or manually creating new, specific templates. Independently of the template used, cell divisions must always give rise to two daughter cells. *SIMI BioCell* stores the data in two text-based file types: an SBC file and an SBD file, the former containing metadata, the latter containing the actual lineage data.

*SIMI BioCell* was with no doubt a pioneer achievement and valuable tool in the field of cell lineaging at the time of its release. It played an important role e.g. in comprehending the cell genealogy of *C. elegans* (Schnabel et al. 1997). However, there are a number of shortcomings for users today. The software is expensive. *SIMI BioCell* was developed for Microsoft Windows XP and only recently updated to run under Microsoft Windows 7. There is no support for the most up-to-date Window systems, or for other major operating systems such as MacOS or Linux. The lineage data can be analyzed only by the tools that are implemented in *SIMI BioCell*, which are limited. Deeper analyses such as the measurement of proliferation, cell division rates, or patterns of cell migration cannot be done. Also, the produced lineage data cannot be exported for analysis in external software (e.g. MATLAB). Furthermore, *SIMI BioCell* can only work properly with data sets that do not exceed a certain size (<50GB in our experience). For data sets that are generated by modern microscopic systems (e.g. light-sheet microscopy) *SIMI BioCell* is not suitable.

To address the issue of visualization and cell lineage reconstruction using terabyte sized datasets generated by light-sheet microscopy, the software *Massive Multi-view Tracker (MaMuT)* was introduced recently (Wolff et al., 2018). It is a plugin for the open scientific image analysis environment *Fiji (Fiji is just ImageJ)* (Schindelin et al., 2012) and builds upon the already existing plugins *TrackMate* (Tinevez et al., 2017), *BigDataViewer* (Pietzsch et al., 2015), and *3D Viewer* (Schmid et al., 2010).

Vellutini et al. (Vellutini et al., 2017) developed a Python library which reads *SIMI BioCell* lineage data for quantitative analysis. They also provided a program called *simi2mamut*, which, building on the library, converts the lineage data to the *MaMuT*-compatible XML format (both available on https://github.com/nelas/simi.py). The authors used these tools successfully to visualize their *SIMI BioCell* data in *MaMuT*. However, both the library and *simi2mamut* had trouble running on many of our own data sets. Also, *simi2mamut* does not take into account meta-data, like spot color and size, which can also be displayed in *MaMuT*.

Here we introduce *BioCell2XML*, a tool that converts *SIMI BioCell* lineage data in a way that (i) the conversion works for a broad set of *SIMI BioCell* project files, (ii) the resulting *MaMuT* lineage, when displayed, represents the original as close as possible (including spot colors and sizes), and (iii), building on user friendly instructions, facilitating it’s use also for non-programmers, thus making a quick transfer from *SIMI BioCell* to *MaMuT* possible for a broad range of scientists.

## Methods

### Image pre-processing

The annotation tool *MaMuT* relies on the file format of the *BigDataViewer* which is the container file format HDF5. In order to be able to use *MaMuT* users need to transfer their image series to this format. *SIMI BioCell* was initially designed for handling the data coming from the 4D microscope developed by Ralf Schnabel (Schnabel et al., 1997). The image format that is set as default in *SIMI BioCell* to store 4D image series is LURAWAVE (LWF, developed by Luratech/Foxit). LWF images first need to be converted into TIFF (tagged image file) format (see work flow in Fig. 1). In a following step the image series can then be converted to H5/XML format, using the *Fiji* platform (either with plugins *BigDataViewer* or *Multiview Reconstruction* (Preibisch et al. 2014)). More detailed instructions for all steps are provided in the supplement.

**Fig 1.**
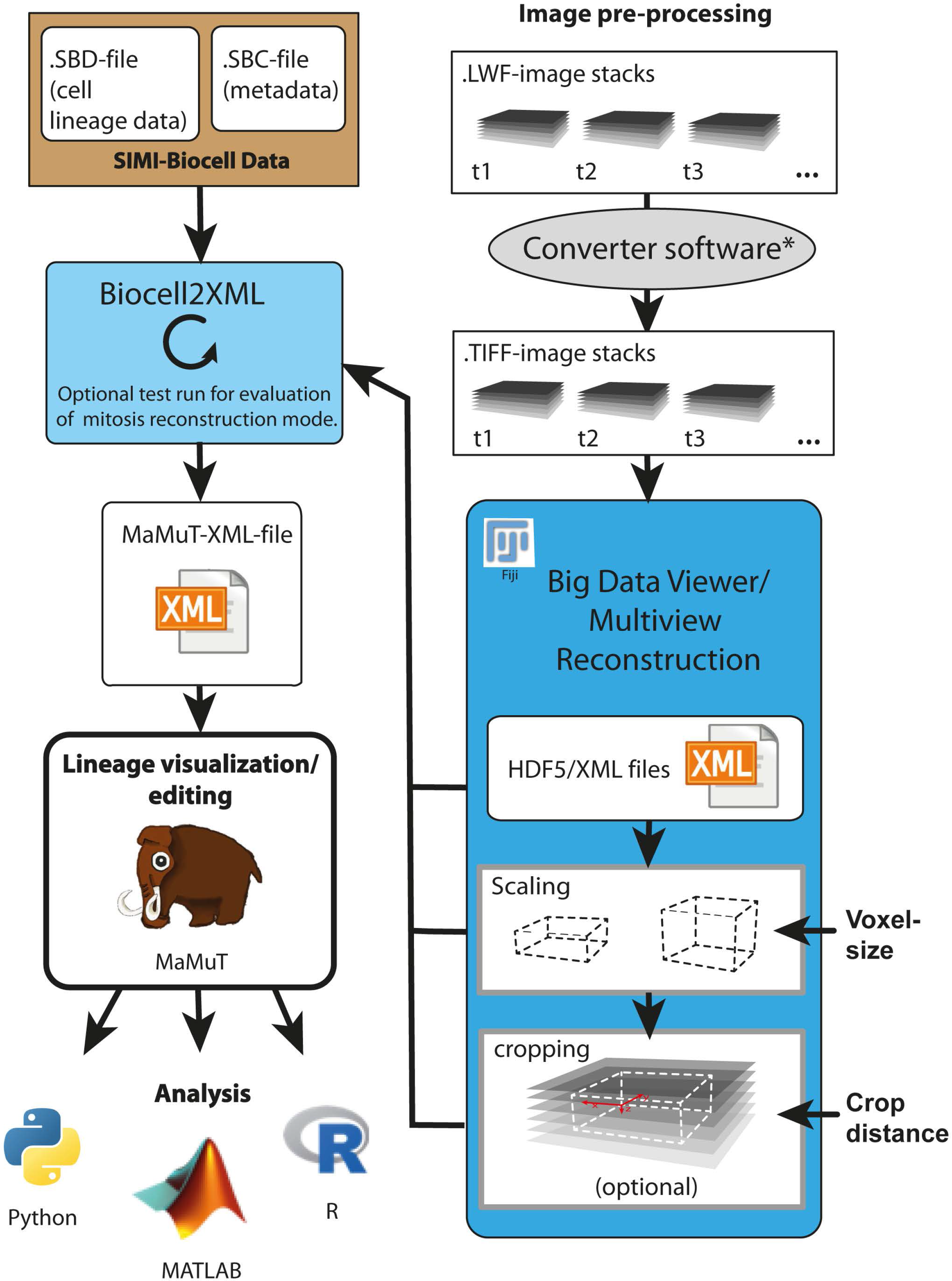
Work flow of data conversion from *SIMI BioCell* to *MaMuT*. The schematic shows the principal work flow of the conversion. On the right side, image pre-processing is depicted. Image data conversion (upper right) is optional, if images are already in TIFF format. TIFF images are converted to H5/XML in *Fiji*. Scaling and cropping can be applied to this data. The H5/XML files, SBD file, SBC file and optionally cropping information and voxel size are required for *BioCell2XML*. Test runs can be performed, and conversions repeated. The output MaMuT/XML file can be used for visualization or various other methods of analysis.

### Running BioCell2XML

*BioCell2XML.py* and a detailed instruction for its use can be downloaded from https://github.com/guleom/BioCell2XML. The software is written in *Python 3* (Python Software Foundation) and will run under most operating systems. *Python 3* and instructions for installation are available at http://www.python.org/.

*BioCell2XML* requires information from three input files: the SBC and the SBD file of the *SIMI BioCell* project, and the H5/XML file which was generated in the image pre-processing step. Note, that the names of all input files and directories must not contain any space or non-ACSII characters.

The SBC, SBD and H5/XML files should be located in the working directory together with the program file BioCell2XML.py. The program is launched by entering

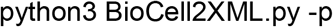

This will initiate a step-by-step input mode where the user is prompted to enter the name of the *SIMI BioCell* project and of the H5/XML file. Subsequently, the software prompts the user to specify all further conversion options, which are explained in detail below (cropping, performing a test run, choice between modes of mitoses reconstruction, transfer of spot metadata, and spot interpolation). The program will then create a *MaMuT* readable XML file with the translated lineage data in the working directory and, if specified by the user, a legend file containing metadata that cannot be displayed in *MaMuT* (like user comments) in TXT format.

Advanced users can run the program non-interactively (useful e.g. for processing numerous lineages) by entering

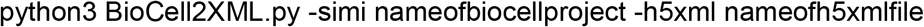

followed by the optional arguments for specific translation settings (see Table 1). This mode also allows importing *SIMI BioCell* and H5/XML files from other than the current working directory.

**Table 1.**
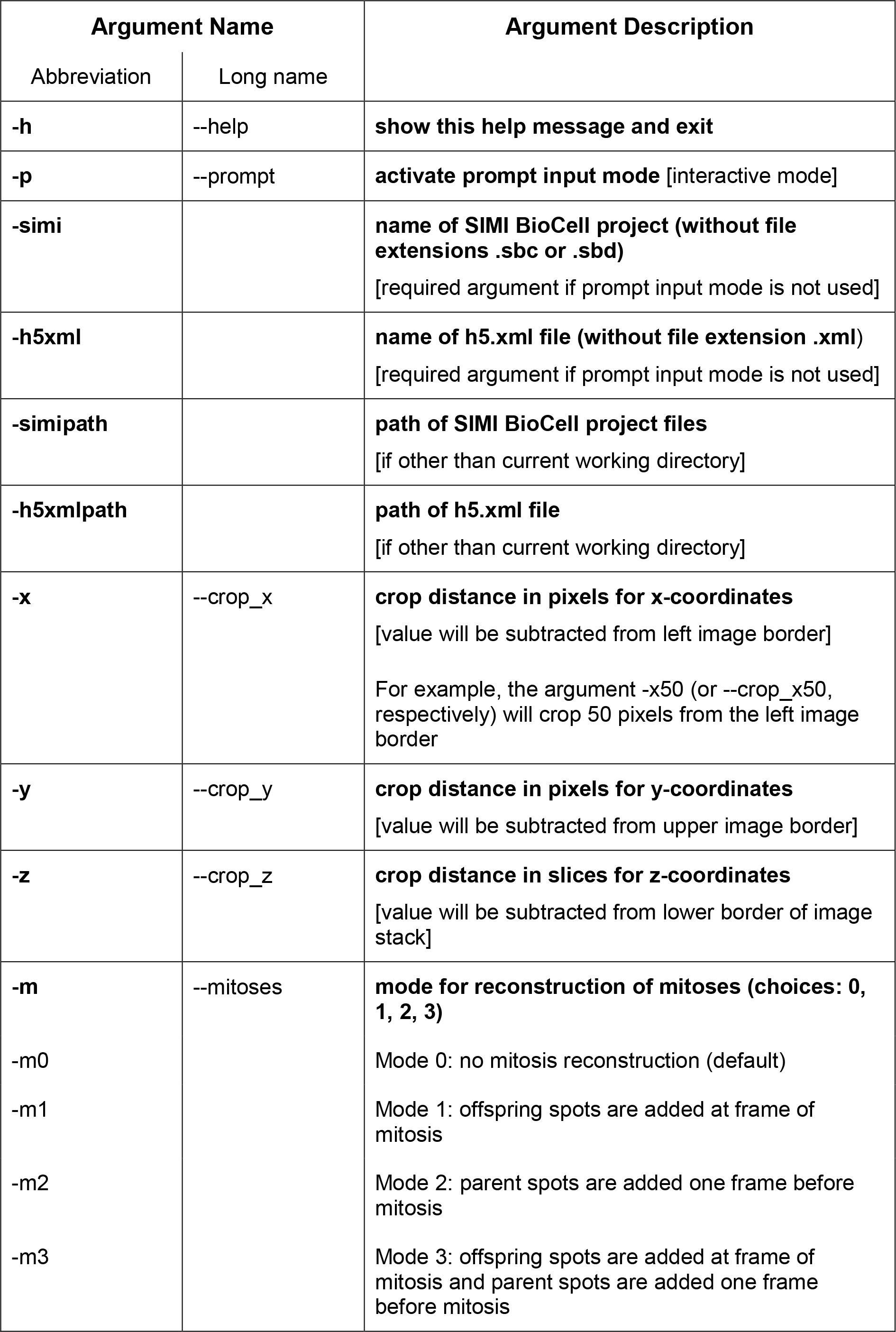
List of arguments for using *BioCell2XML.py* in command-line interpreter mode.

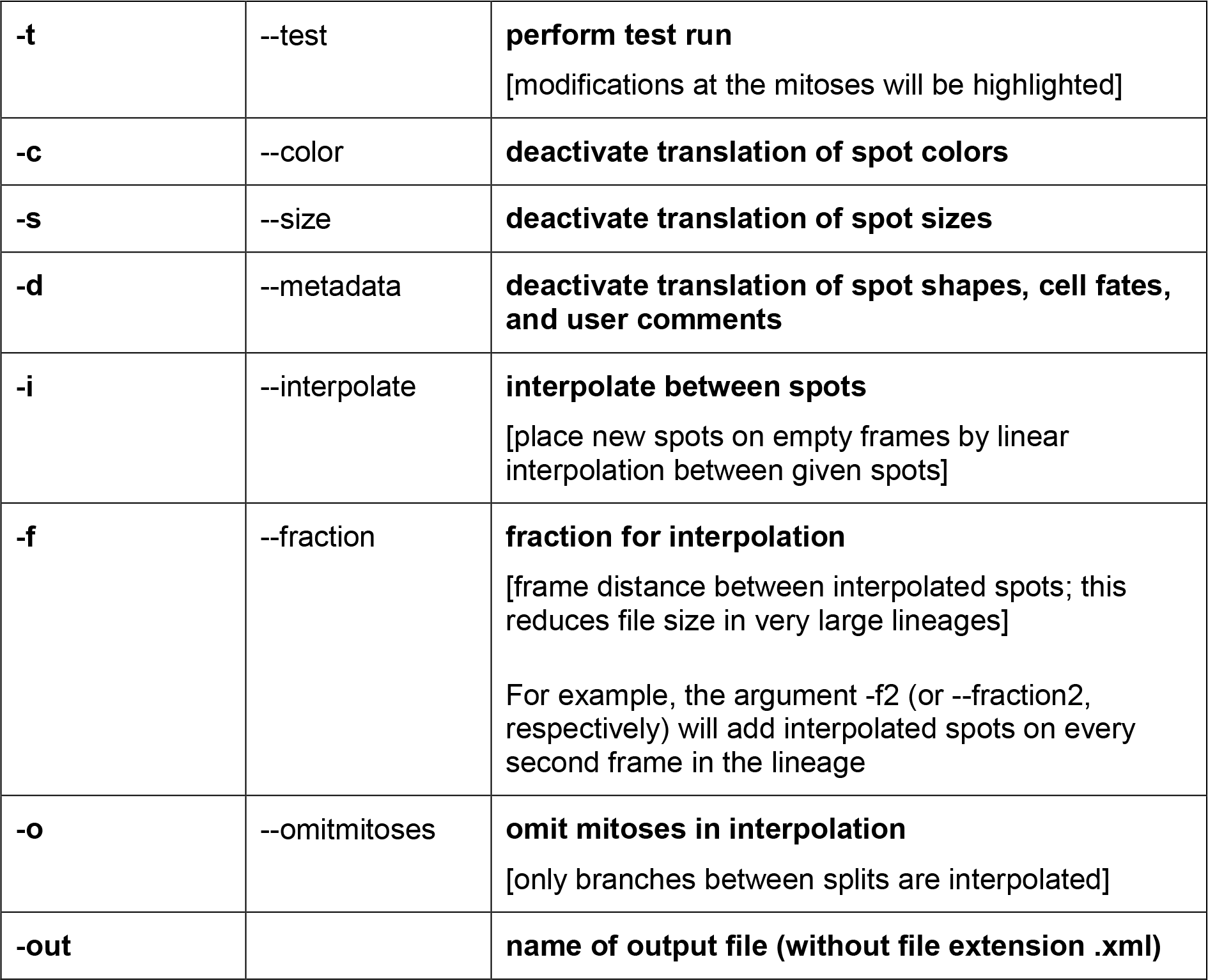

## Results

### Basic functionality of BioCell2XML

*BioCell2XML* traverses the *SIMI BioCell* project files and extracts scan setting from the SBC and cell lineage data from the SBD file. Scaling information (image resolution, voxel size, number of frames and slices) is read from the H5/XML file. For all active cells in the SBD, name, mitosis frame, fate, size, shape, color, user-specified comments, and all spots with timeframes and x/y/z coordinates are collected. The data are updated with time and resolution information from the SBC and H5/XML files, and converted to the spot format used in *MaMuT* XML files. In a next step, the genealogy of the cell lineage is recovered from the order of cells in the SBD file, and this information is translated to the XML format, which uses tracks and edges (connections of consecutive source and target spots). If specified by the user, in further steps, first, cell divisions and mitoses spots are reconstructed and/or, second, new spots are added by interpolation. Finally, all spot, track and edge data, and scaling information is written to the output XML file and saved to the working directory. An additional legend TXT file is outputted, if the *SIMI BioCell* lineage includes alternative branches or mitoses that have only one daughter cell. If the *SIMI BioCell* lineage contains metadata and user comments, which cannot be displayed directly in *MaMuT*, these are written the XML and the legend file, if the user wishes to.

### Cropping

Images in LWF format have a reduced file size thanks to the used compression technology. When the images are converted to TIFF format, the data size will increase approximately 10-fold and reduced again 2-fold when converted to compressed H5/XML. Particularly for large data sets it is advantageous to crop empty regions from the images before conversion. Cropping can be performed by cropping a hyperstack in *Fiji* or batch cropping in any other software. The cropped images can be opened as hyperstack in *Fiji*, and then exported as H5/XML format using the plugin *BigDataViewer*. Detailed information on how to perform this cropping is given in the supplementary file *BioCell2XML_User_Guide.pdf*. Cropping of the image data demands correction of the spot coordinates in the lineage data, in order to maintain the correct spot positions. Therefore, users should note the new x, y and z values that are defined for cropping (Fig. 1). This data has to be entered when running *BioCell2XML*.

### Reconstructing cell divisions

Mitosis is a continuous process, which can only be accounted for to a limited extent by recording discrete events (Fig. 2A). *SIMI BioCell* and *MaMuT* apply different strategies to represent cell divisions in the lineage data.

**Fig 2.**
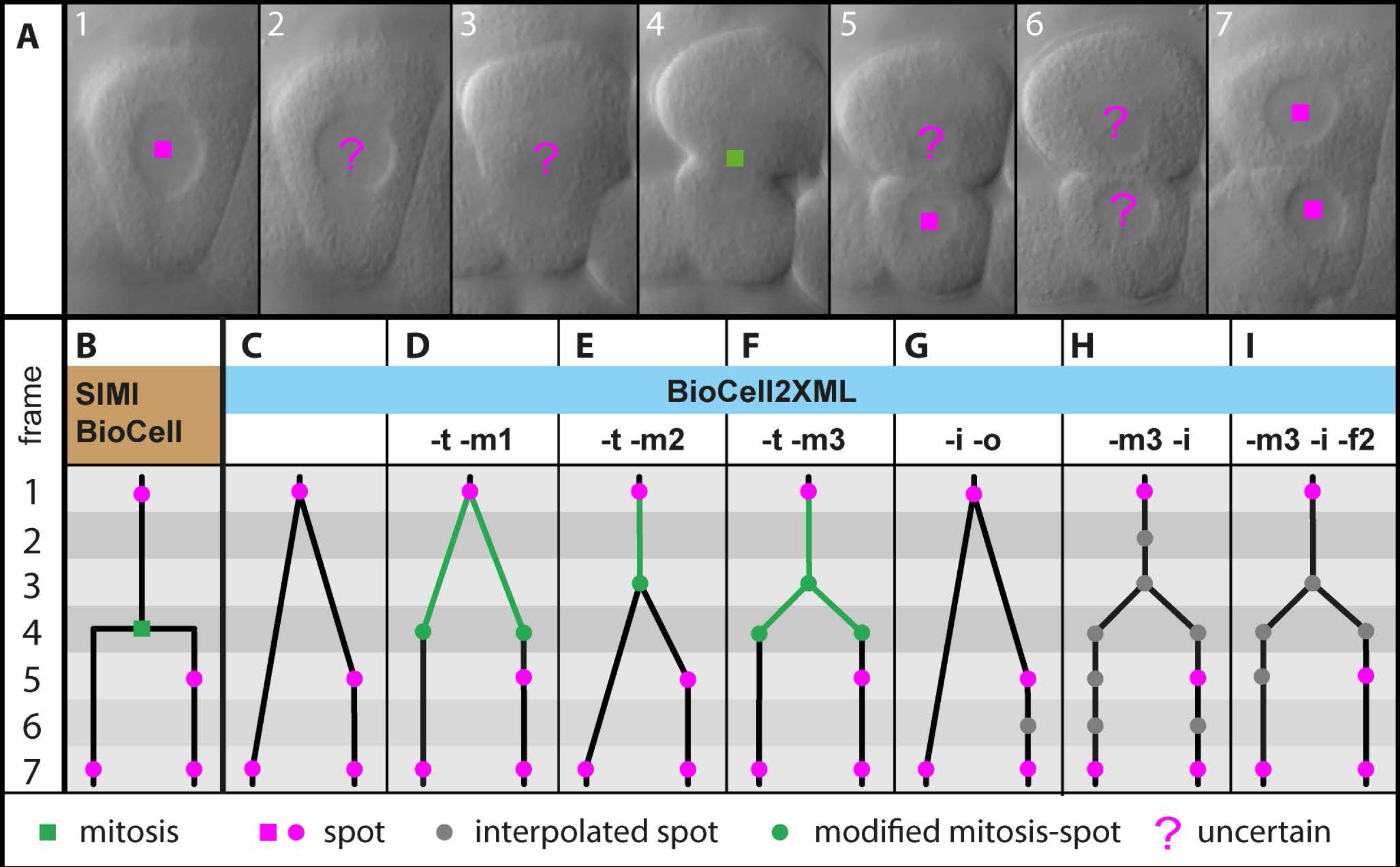
Modes of mitosis reconstruction and interpolation. **A.**Example image array covering seven frames, showing a lineaged blastomere of *Psammechinus* sp. Cells are marked with spots only where nuclei are distinctly visible. A mitosis was inserted in frame 4. **B-I**. Different possible lineaging approaches based on the example in **A**. The line above **C-I**shows the argument combinations for *BioCell2XML* for the respective mitosis reconstruction mode (see Table 1). **B**. Lineage representation as in *SIMI BioCell*. **C**. *SIMI BioCell* lineage converted to *MaMuT* following default settings. **D**. Converted lineage with reconstructed daughter spots. **E**. Converted lineage with reconstructed mother spot. **F**. Converted lineage with both, reconstructed daughter spots and reconstructed mother spot. **G**. Lineage with interpolation only outside mitosis. **H**. Interpolated lineage with default interpolation setting. **I**. Interpolated lineage with fraction = 2.

In *SIMI BioCell*, splits in the lineage are added independently of placing spots. The last spot has to be placed at least one frame before the split, while the daughter spots can be placed at the mitosis frame or later. Adding the split event in *SIMI BioCell* records the mitosis time and introduces two new branches on which spots of the two daughter cells can be placed. *MaMuT*’s philosophy of representing mitoses is stricter. Here a mitosis can be represented only by edges, connecting a mother spot with two daughter spots. As in *SIMI BioCell* the last mother spot and the first daughter spot have to be separated by at least one frame.

Lineaging strategies vary between users and depending on the appearance of the investigated embryos. E.g. scientists working on species with distinctly visible nuclei will place the spots at the position of the nucleus when lineaging (e.g. frame 1 and 7 in Fig. 2A). Nuclei disintegrate during mitosis, often to a degree that makes them indistinguishable from the surrounding cytoplasm. Investigators will then either choose vague spot positions or leave a few frames empty (e.g. frames 2-6 in Fig. 2A). In a direct translation of a *SIMI BioCell* lineage to *MaMuT*, as performed by default, empty frames before the mitosis frame will lead to the split appearing earlier than initially intended by the user (compare Fig. 2B and C). Depending on the type of analysis that is to be performed on the lineage, this can be unwanted, e.g. if quantitative analyses of cell division times is to be performed. For this reason, *BioCell2XML* allows different modes of mitosis reconstruction. Users can choose to create additional daughter spots at the mitosis frame (Fig. 2D). The additional spot is generated by linear interpolation between the position of the last mother spot and the position of the first daughter spot onto the frame of mitosis. Similarly, an interpolated mother spot can be added to the lineage one frame before the mitosis frame (Fig. 2E). Finally, both these additional spots can be reconstructed (Fig. 2F).

In many cases it may be difficult for users to decide how strong the impact of a chosen mitosis reconstruction mode will be. Therefore, *BioCell2XML* provides the option of a test run. If selected, this option outputs a *MaMuT* lineage where the affected spots and the edges leading to their mother spots, are colored against a grey background. Thereby, the green color marks spots that were added by the mitosis reconstruction algorithm, while spots that were present in the original *SIMI BioCell* lineage and that coincide in timeframe with a mitosis reconstructed spot are marked in magenta. Users can use this option to visually check the impact of the available reconstruction modes and decide how to proceed.

Another obstacle occurs when *SIMI BioCell* lineages contain mitosis events, which have only one daughter cell. Single-daughter mitoses can be the result of limited visibility of one daughter cell in the microscopic images, or of incomplete lineaging. Either way such a mitosis would be displayed in *MaMuT* as a continuous branch of the lineage tree. Users can identify these hidden splits by using the test run option in combination with any mitosis reconstruction mode. In the resulting lineage, the last mother spot and/or the first daughter spot (depending on the chosen reconstruction mode) can be recognized as colored spot within a continuous branch.

*SIMI BioCell* also offers the possibility to add so-called alternative branches inside a lineage. Users may employ this option, when they are uncertain about the correct cell track. *BioCell2XML* will translate such alternative branches as polytomous mitosis event. Since polytomies are not allowed in *SIMI BioCell*, the presence of a polytomous mitoses in the translated *MaMuT* lineage points to the presence of an alternative branch in the original *SIMI BioCell* data. Since mitoses with only one daughter cell as well as alternative branches are lineage events that the user may revise and edit in *MaMuT* later, *BioCell2XML* will list the mother spots of these events in the legend TXT file.

### Spot sizes

In *SIMI BioCell*’s 3D View window, users can set spot sizes to “small”, “medium” or “large”. The spots can then be scaled all together. By default, “small” and “large” spots are displayed at approximately 0.05 and 0.07 times the width of the displayed volume, respectively. These proportions can vary somewhat when the view is rotated. *BioCell2XML* can translate the spot sizes from the SBD file to size values that give similar size differences when displayed in *MaMuT*’s 3D Viewer. If spot sizes were not set in *SIMI BioCell*, the default spot size of *MaMuT* will be used (20 pixels, referring to the resolution of the non-scaled images). In any case, if in *MaMuT* spots are added manually to a translated *SIMI BioCell* lineage, the new spots will always have *MaMuT*’s default spot size.

### Colors

Manually specified colors in *SIMI BioCell* lineage data are given as single numbers in OLE-code. If R, G and B represent red, green, and blue values (0 – 255) in an 8-bit color space, the OLE value for this color is defined as

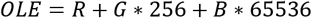

*MaMuT* uses a different but similar method to code RGB colors, which we refer to as WIN here. It can be described as

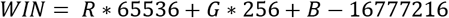

Using this information, *BioCell2XML* will calculate the corresponding color value for *MaMuT* from the spot colors given in the SBD file.

### Interpolation

For various reasons it can be desirable to create interpolated spots between recorded spots in the lineage. Different types of analyses may require spot information for each frame. Also, visualization of development over time in *MaMuT*’s 3D Viewer will be more convenient if lineages are interpolated. Unlike *SIMI BioCell*’s 3D View this tool does not interpolate spots for visualization on the fly. *MaMuT* provides a built-in interpolation option but it will not transfer manually set spot colors to interpolated spots. Therefore, we included an option with *BioCell2XML* that adds interpolated spots, which are calculated by linear interpolation between the positions and sizes of the original spots. If the color of source and target spot should differ, interpolated spots will adopt the source spot color for the first half of the frame distance, and the target spot color for the second half.

Interpolation can interfere with the desired strategy of translating mitoses, e.g. if users wish to translate the lineage using the default mitosis reconstruction to then edit the mitosis events manually in *MaMuT*. For this case, *BioCell2XML* provides the option to omit the mitoses in the interpolation (Fig. 2G). Interpolation can otherwise be performed together with any mode of mitosis reconstruction (e.g., mode 3, Fig. 2H).

The interpolation option must be handled with care for large lineages, because file size can increase significantly through the newly introduced spots. For this reason, *BioCell2XML* includes an additional option for interpolation, in which the interpolation fraction can be set. A fraction of two will introduce new spots only for every second frame in the lineage, if this is not in conflict with the chosen mitosis reconstruction mode (Fig. 2I).

### Comments and other metadata

Part of the metadata saved in *SIMI BioCell* cannot be displayed directly in *MaMuT*. This is true for the displayed spot shape (the semi-schematic spot representation as cube, diamond, etc. used in *SIMI BioCell*’s 3D View), cell fate information, and user-generated comments. We consider it important, nevertheless, to allow users to keep this information in the file, so it can be accessed later, e.g. using other software. Unless specified otherwise by the user, *BioCell2XML* will write the non-*MaMuT* compatible metadata into a special section of the XML file. Metadata that are float or integer numbers are displayed in the spot information window of *MaMuT*’s TrackScheme. Unfortunately, this will not work for string data, like user comments. *BioCell2XML* will therefore create a legend text file, which can be opened with any text editor and where these metadata are listed for the respective spots.

## Examples

*BioCell2XML* allows users to transfer *SIMI BioCell* lineage data to the *MaMuT* readable XML format. Fig. 3 shows an example cell lineage displayed in *SIMI BioCell* (Fig. 3A-D) and the converted lineage in *MaMuT* (Fig. 3E-H). Both programs provide the same basic features: 3D visualization (Fig. 3A, E), cell tracking window showing the image data (Fig. 3B, F), overview of the lineage tree (Fig. 3C, G), and detail of the lineage tree (Fig. 3D, H). Using default settings results in a 3D representation (Fig. 3E) that differs little from the appearance in *SIMI BioCell* (Fig. 3A). In this example, the image data was cropped, omitting the empty regions around the embryo. Users can take advantages of several *MaMuT* features, like focus-dependent scaling of spots in the *MaMuT* viewer (Fig. 3F), interactive slicing of the image volume, smoother visualization of rotation in the 3D Viewer, opening multiple viewer windows, e.g. showing x, y and z view of the volume simultaneously, or conveniently editing the lineage in the tree window, e.g. deleting or reconnecting spots and edges.

**Fig 3.**
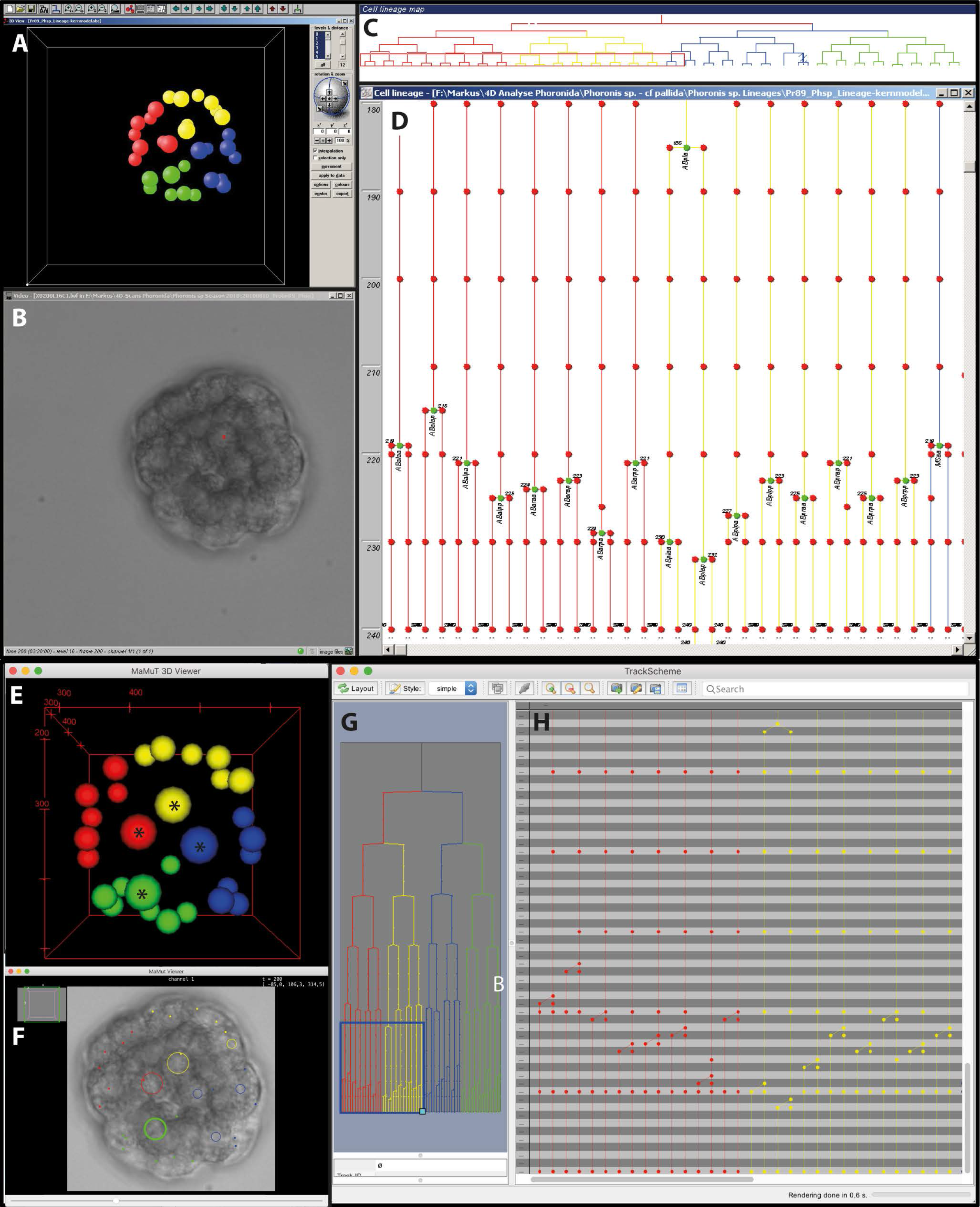
Example for converted lineage data. **A-D**. Different windows of *SIMI BioCell*, displaying an example lineage of *Phoronis* sp. **A**. 3D View. **B**. Cell tracking window. **C**. Overview of lineage tree. The slender red rectangle over the lower left part of the tree marks the section displayed in **D**. **D**. Lineage tree window. **E-H**. Corresponding windows of *MaMuT*. **E**. 3D Viewer. **F**. *MaMuT* viewer. **G**. Overview of lineage tree in TrackScheme window. The blue rectangle over the lower left part of the tree marks the section displayed in **H.** Mitosis reconstruction mode 3 (Fig. 2F) was used. **H**. Detail of lineage tree in TrackScheme window. Arrows in **A**and **E** mark the spots for which the size was set to “large” in *SIMI BioCell*. The remaining spot sizes were set to “small”.

Image data can be scaled to actual voxel size (not provided in *SIMI BioCell*), which facilitates geometric analyses of the data. *MaMuT* also does not restrict divisions to being dichotomous. In the current version, spots can have two or more daughter spots, which could be convenient for some tracking projects not restricted to cell division. Cell fusion, which occurs e.g. in muscle development, can be documented with *MaMuT* as linking events between different lineages. *MaMuT* does not provide the same options for showing labels as *SIMI BioCell*. Spots can be named individually, but cells cannot, as cells are not defined as objects in the lineage data. *BioCell2XML* nevertheless provides access to and usability of this metadata.

Analyses that can be applied to converted *SIMI BioCell* lineages are manifold. We present two basic examples here. Fig. 4A-F shows converted lineage data of *Psammechinus* sp. (kindly provided by Thomas Stach). The figure demonstrates that optical sections of DIC-image stacks in different dimensions (Fig. 4A-C) can improve orientation in the 3D space.

**Fig 4.**
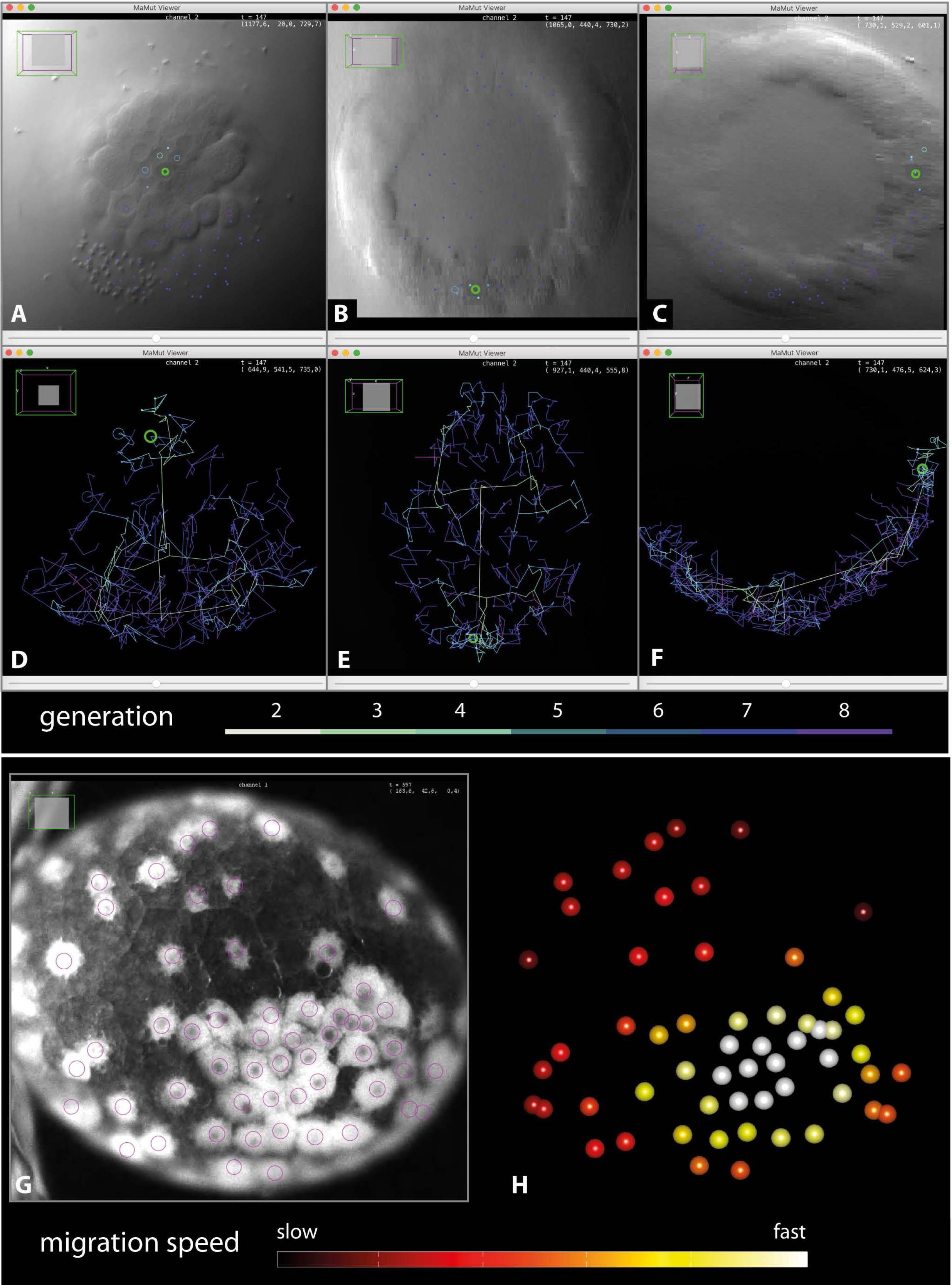
Examples for post-conversion analyses. **A-F**. Example of multiview display and analysis of a converted lineage of *Psammechinus* sp. in *MaMuT*. **A-C**. Optical sections of embryo in z, y and x view, respectively. A highlighted green circle marks the same spot in all views. **D-F**. Plotted cell tracks for same embryo in the same orientations as in **A-C**. Tracks were color coded for cell generations. **G**, **H**. Example view of a cell lineage of *Cryptorchestia garbini* (syn. *Orchestia cavimana*, original data from Hunnekuhl & Wolff, 2012). **G**. Embryo during germ disc condensation. Circles mark the cells that are annotated in *SIMI BioCell*. **H**. Plot created in MATLAB, showing the annotated cells of the embryo in **G** but color coded for their migration speed (from slow: dark to fast: light).

In addition, the cell tracks are given a color code that changes gradually with each division cycle, by subsequently using two Python scripts (*lineage_from_mamut*, and *set_colors*, available at https://github.com/jirikowg/Pytools_4_MaMuT) (Fig. 4D-F). A converted cell lineage of the amphipod crustacean *Cryptorchestia garbini* (syn. *Orchestia cavimana*) (Fig. 4G, original data shown in Hunnekuhl & Wolff, 2012) was processed in MATLAB (Mathworks, USA) to compare speed (color coded) for cells that condense into an early germ disc (Fig. 4H).

The different methodologies of lineage representation applied by *MaMuT* and *SIMI BioCell* demand that users consider briefly their personal lineaging strategy and the goal of the conversion to *MaMuT*. *BioCell2XML* allows both, conversion with priority for visualization, thus giving a result that most closely represents the lineage displayed in *SIMI BioCell*, as well as conversion for further editing and quantitative analyses.

## Acknowledgements

We are grateful to Tim Strickler for helping with color conversion from OLE code, to Thomas Stach for providing example data sets of *Psammechinus* sp. for this study, and to Bruno Vellutini for assisting with the use of *simi2mamut*. We also would like to thank Bruno Vellutini for helpful comments on the manuscript.

